# Is there such a thing as landscape genetics?

**DOI:** 10.1101/018192

**Authors:** Rodney J. Dyer

## Abstract

For a scientific discipline to be interdisciplinary it must satisfy two conditions; it must consist of contributions from at least two existing disciplines and it must be able to provide insights, through this interaction, that neither progenitor discipline could address. In this paper, I examine the complete body of peer-reviewed literature self-identified as landscape genetics using the statistical approaches of text mining and natural language processing. The goal here is to quantify the kinds of questions being addressed in landscape genetic studies, the ways in which questions are evaluated mechanistically, and how they are differentiated from the progenitor disciplines of landscape ecology and population genetics. I then circumscribe the main factions within published landscape genetic papers examining the extent to which emergent questions are being addressed and highlighting a deep bifurcation between existing individual- and population-based approaches. I close by providing some suggestions on where theoretical and analytical work is needed if landscape genetics is to serve as a real bridge connecting evolution and ecology *sensu lato*.

## Introduction

As the landscape of scientific research and new discovery becomes increasingly interdisciplinary, in both the makeup of diffuse research teams and in the composite nature of hypotheses being examined, it is becoming important to quantify the way in which individual disciplines interact and how new ones contribute to knowledge generation. According to a National Academies report entitled, *Facilitating Interdisciplinary Research* (2004), interdisciplinary research is “...a mode of research by teams or individuals that integrates information, data, techniques, tools, perspectives, concepts, and/or theories from two or more disciplines or bodies of specialized knowledge...” The purpose of these interactions are to either “...advance fundamental understanding or to solve problems whose solutions are beyond the scope of a single discipline or area of research practice.” These specific outcome-orientated restrictions delineate interdisciplinary research from mere collaborations and/or aggregated multidisciplinary work, which may satisfy the composition of the research teams though not fulfill the stated purposes in providing emergent insights. These restrictions beg the question of whether research we are currently conducting and referring to as interdisciplinary is actually addressing emergent questions (heretofore unanswerable using normal approaches from constituent disciplines) or are we simply borrowing approaches and techniques established from different disciplines to answer the same research questions we have been examining for some time.

As an example, this manuscript looks at a body of research self-identifying as *landscape genetics*, an emerging keyword designation being applied to research projects studying how ecological, vegetation, anthropogenic, and topographic context influence genetic connectivity and structure in both natural and modified populations. The moniker was originally introduced by Manel *et al.* (2003), who provided a definition based upon mechanisms, “The two key steps of landscape genetics are the detection of genetic discontinuities and the correlation of these discontinuities with landscape and environmental features such as barriers.” While the mechanics of barrier identification and landscape correlation are not to population genetics (e.g., Dobzhansky 1948; Merriam *et al.* 1989, Keyghobadi *et al.* 1999, etc.), the ubiquity of model-based clustering approaches initiated by Pritchard *et al.* (2000) and the advancements in spatial ecological analysis and GIS technology set the stage for this potentially interdisciplinary research approach. Because of this mix using spatio-ecological data to predict genetic characteristics of organisms (now much broader than just discontinuities), landscape genetics has been characterized as an interdisciplinary *fusion* of landscape ecology and population genetics (e.g., Manel *et al.* 2003, Storfer *et al.* 2006, Holderegger & Wagner 2008, Balkenhol *et al.* 2009). Given the focus of these constituent fields and the kinds of research questions being classified as landscape genetics, it is an obvious supposition. However, if landscape genetics is to be correctly defined as an interdisciplinary field, it needs to be more than just the rough interdigitation of progenitor disciplines. Moreover, it must be able to provide insights into processes that are heretofore unattainable—failing to do so would suggest that its continued use would only add an arbitrary and somewhat meaningless categorization to our research descriptions.

Its flagship journal, *Landscape Ecology*, defines the discipline itself as an “...interdisciplinary science that focuses explicitly on the ecological understanding of spatial heterogeneity”, providing a bit of recursive irony to the definition of landscape genetics. In the preface to the edited volume, *Landscape Ecology: A Top-Down Approach*, Sanderson & Harris (2000) suggest the main component that makes landscape ecology distinct from other fields of ecology is that “...it explicitly encompasses and builds upon the role of heterogeneity in space as well as time.” It can be argued that the primary role that landscape ecology has played in the formation of landscape genetics is through analytical advances and integration of sophisticated approaches to quantifying spatial heterogeneity, particularly through the integration of geographic information systems (GIS) into their analyses. In most landscape genetic studies, these techniques provide the basis for the predictor set of variables under consideration. Despite the specifics of the contribution, there is a clear point at which a landscape ecological study will be categorized as landscape genetics; namely when the study includes the use of genetic markers.

Identifying the characteristics of hypotheses that separate landscape genetic studies from population genetic ones is much more difficult. Due in part to its longevity, population genetics has developed a more broad scope of research directions, focusing on both micro- and macro-evolutionary mechanisms. Population genetics, in both theory and practice, examines the mechanisms of evolutionary processes and how they influence within and among groups dynamics. There are clearly kinds of research questions in population genetics that would not be easily mistaken for landscape genetics, as they do not include the environment through which populations are interacting. However, there is a long history of population genetic studies that have specifically included ecological and/or spatial data in the analyses of genetic connectivity and structure (e.g., Smouse *et al.* 1987, Piertney *et al.* 1998, Pannell & Charlesworth 1999) making large fractions of population and landscape genetic studies virtually indistinguishable. As such, the inclusion of spatio-ecological predictor variables alone does not reclassify a population genetic study as landscape genetics and identifying the boundaries separating population from landscape genetics much less clear.

Applying model-based clustering of genetic information with external landscape and ecological features may not be sufficient to justify the use of interdisciplinary or to support the *de novo* creation of a sub-discipline. For that, the outcome of the research must advance fundamental understandings or provide insights that progenitor disciplines could not ascertain. Looking back at the definition, Manel *et al.* suggest this very thing saying, “Landscape genetics can resolve population substructure across different geographic scales at fine taxonomic levels, thus it is different from the existing understanding of the microevolutionary processes that generate genetic structure across space.” At a minimum, this suggests that landscape genetics defines a novel body of knowledge able to identify emergent processes that either landscape ecology or population genetics could not characterize. These processes used in landscape genetics can specifically address the causative ecological and spatial forces influencing the formation and maintenance of genetic structure in a way that was previously unreachable. If true, then landscape genetics is indeed interdisciplinary and it may contain insights that, through introgression of either methodological or theoretical approaches, would benefit both landscape ecology and population genetics greatly. More importantly though, if landscape genetic studies do define a cohesive set of hypotheses and/or approaches that are divergent, then it has the potential to serve as a direct conduit through which fundamental understandings within both Ecology or Evolution (*sensu lato*) may be exchanged bringing the evolutionary process into tighter connection to the ecological contexts within which they operate. A lofty goal indeed!

Here I examine the body of literature defined as landscape genetics to determine if it contains the two components—contributions from two or more disciplines and emergent insights—necessary for it to be considered interdisciplinary research. Rather than summarizing the results from recently published empirical papers (there are already enough review papers on this topic), I instead examine the textual structure of all manuscripts self-identified as landscape genetics using methods of text mining and natural language processing. From these manuscripts and a representative sample of ones unambiguously classified as landscape ecology and population genetics, I quantify the composition of the *Introduction* and *Methods* sections to determine if the kinds of questions being asked and the ways in which the research is implemented support landscape genetics as an interdisciplinary field addressing emergent hypotheses. In light of the results generated, I discuss the current state of landscape genetics, its relationship to its progenitor disciplines, and provide suggestions for future research directions that aid in understand the interaction of ecological context, spatial heterogeneity, and microevolutionary processes. Moreover, I hope the results presented spark continued development of both theory and applications for this unique intersection of ecological and evolutionary research.

## Methods

### Manuscript Collection

Data were collected from literature searches conducted using ISI Web of Science in July 2014. The body of literature designated as landscape genetic (hereafter LG) consisted of the manuscripts that both cited the original Manel *et al.* (2003) manuscript and specifically self-identified by inclusion of the words “landscape genetic” as a component of the supplied keywords, the title, or contained within the *Abstract* (*n.b.*, the plural was also used). Similar collections of manuscripts self-identified as landscape ecology (LE) and population genetics (PG) spanning the same time period (2003–2014) were collected. Since the number of LE and PG manuscripts published outnumber LG papers by an order of magnitude, a random sample of 100 manuscripts were selected from each group. Not all potential manuscripts were available for inclusion, PDF documents from the journals *Ecoscience, Israeli Journal of Ecology and Evolution, Invertebrate Systematics*, and the *International Journal of Sustainable Development & World Ecology* were behind paywalls and were unavailable to my institution. Manuscripts from *Theoretical Population Biology* and the *American Journal of Physical Anthropology*, while available, contained non-conforming PDF encodings making the content unparsable by the software written for this work.

Raw text content was preprocessed and prepared for statistical analysis in R (R Core Team, 2014) as follows. All PDFs were converted into ascii text format using the footprints library (Dyer, unpublished; available from http://dyerlab.github.com/footprint). Errors in pdf to text conversion (e.g., the loss of a white space separating words, non-ascii ligature translation, etc.) were hand checked and corrected as necessary. Next, the raw text was extracted for both the *Introduction* and *Methods* sections separately; to differentiate between the way a manuscript formulates the research question from how the research is implemented. It is implicitly assumed that content in the *Results* section would be largely redundant to what is presented in the *Methods* and that the *Discussion* is where the context of the findings are made, not where key features that differentiating LG from PE and PG are found. All LG papers were classified by hand using four potentially overlapping categories (Table 1): *Review, Simulation, Animal*, and *Plant.* Each manuscript could have more than one categorization but no manuscript was left uncharacterized. Only manuscripts determined to be empirical in nature (e.g., those that are not classified as *Review*) were used in the following analyses. Both LE and PG manuscripts determined as *Review* were similarly rejected.

**Table 1.** Topic categories used to classify all self-identified landscape genetic manuscripts. Each manuscript was ascribed at least one of these categories.

| ***Category*** | ***Criteria*** |
| --- | --- |
| *Review* | The manuscript contained a review of existing landscape genetic studies and/or how landscape genetics could be leveraged into an existing field of study |
| *Simulation* | The manuscript either conducted simulations and/or modeling as a component of the work or introduced novel statistical approach requiring the development of new approaches |
| *Animal* | The manuscript analyzed an empirical data set from an animal species |
| *Plant* | The manuscript analyzed a data set from a plant, algal, and/or fungal species. |

The textual structure was preprocessed using the *tm* R package (Feinerer *et al.* 2008, Feinerer & Hornick 2014). Raw text was filtered by removing punctuation, numbers, exogenous white-space, and then converted to lowercase. Common English words (e.g., ‘stop-words’ as defined by Rajaraman & Ullman 2011) were removed using both the “en” and “SMART” libraries (Lewis 2004). Words were then stemmed using the *SnowballC* library (Bouchet-Valat 2014) to retrieve their radicals (e.g., the base English word or word component) preventing differences in word tense and alternative suffixes from artificially inflating the error variance term in subsequent discriminant and cluster analysis. Stemmed text content was translated into a multivariate term frequency vector, 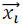. The 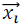 vectors representing the frequency array of stemmed words were then standardized to unit length to minimize bias due to differences in document length as suggested by Dhillon & Modha (2001) for vector space models. Combined across all documents, the raw data are represented as:

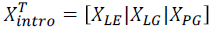

and

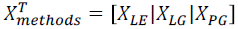

providing a standardized data matrix for each component of the manuscript representing the frequency distribution of word roots.

### Fusion Analysis

To determine the extent to which LG manuscripts are indeed a fusion of LE and PG, a discriminant analysis was performed following Johnson & Wichern (1992) to derive a set of discriminant functions that maximally separates X_*LE*_ and *X*_*PG*_ *Introduction* and *Methods* sections. The discriminant functions were defined on separate covariance estimates for *X*_*LE*_ and *X*_*PG*_, denoted *∑*_*LE*_ and *∑*_*PG*_. Then *X*_*LG*_ manuscripts were classified, assuming *priors* equal to the relative frequency of LE and PG papers (denoted as *p*_*LE*_ and *p*_*PG*_) as belonging to the LE category when

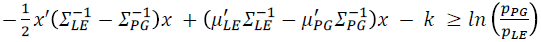

and to the PG category when

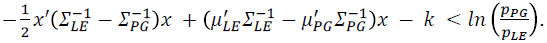

The applicability of derived discriminant functions were evaluated using the assignment error rate (e.g., the mis-assignment of LE or PG manuscripts). Equality of assignment of LG papers to LE and PG discriminant regions indicating an equal contribution of both progenitor discipline to the body of LG manuscripts, was quantified using a binomial test.

### Circumscribing Landscape Genetics

Compositional makeup alone is insufficient to categorize landscape genetics as an interdisciplinary field, it also requires the ability to gain insights that are beyond the scope of the original disciplines. From its inception though, it has been suggested that landscape genetics “...is different from the existing understanding of the microevolutionary processes that generate genetic structure across space.” (Manel *et al.* 2003), satisfying via assertion this requirement. To identify potentially emergent insights that the field of landscape genetics brings, the frequency distribution of stemmed terms in the *Introduction* and *Methods* sections of LG, LE, and PG manuscripts were compared to identify terminologies overrepresented in each discipline. Overrepresentation of specific terms may provide inferences into how landscape genetics provides insights into these microevolutionary processes unappreciated in the other disciplines. Next, the set of self-defined LG manuscripts was examined to determine if they are a single cohesive body or if there are subdivisions within. The vectors in *X*_*LG*_ were translated into pairwise Euclidean distance and a hierarchical clustering was performed using Ward's minimum variance method (R Core team, 2014). Confidence on partitions within the clusters was estimated using a bootstrapping approach (via the pvclust R library, Suzuki & Shimodaira 2011 with 10,000 bootstraps). Differences between major clusters within *X*_*LG*_ were then examined qualitatively by estimation of a term frequency matrix and comparison of the most commonly used terms in the main clusters.

## Results

### Manuscript Collection

The number of manuscripts citing Manel *et al.* (2003) listed in ISI Web of Science (WOS) in July 2014 was 759. From this group, 288 manuscripts were retained that self-identified by using the term ‘landscape genetic(s)’ in the title, abstract, or keywords. A similar search on WOS revealed 3,768 PG manuscripts refined using the term “evolutionary biology.” From these manuscripts 179 were removed as they contained the term “landscape genetic” or “landscape ecology” in the title, abstract, or keywords and thus would not be appropriate for inclusion in discriminant functions describing putatively pure PG papers. Searches for LE manuscripts refined by “ecology” yielded 3,646 manuscripts, of which 259 also contained the search terms “landscape genetics”, “population genetics”, or “genetics” in the title, abstract, or keywords and were similarly removed. A random selection of 100 manuscripts defined as non-review were randomly selected from the LE and PG repositories and used for subsequent analyses. All self-identified LG manuscripts were manually categorized following the definitions in Table 1. By far, the vast majority of LG studies focus on animal systems (*N*_*Animal*_ = 194) with allocation of manuscripts into all of the remaining categories being roughly equal (*N*_*Plant*_ = 61, *N*_*Review*_ = 53, and *N*_*simulation*_ = 52). Nearly a quarter (24%) of the LG manuscripts were assigned more than one category (Figure 1).

**Figure 1.**
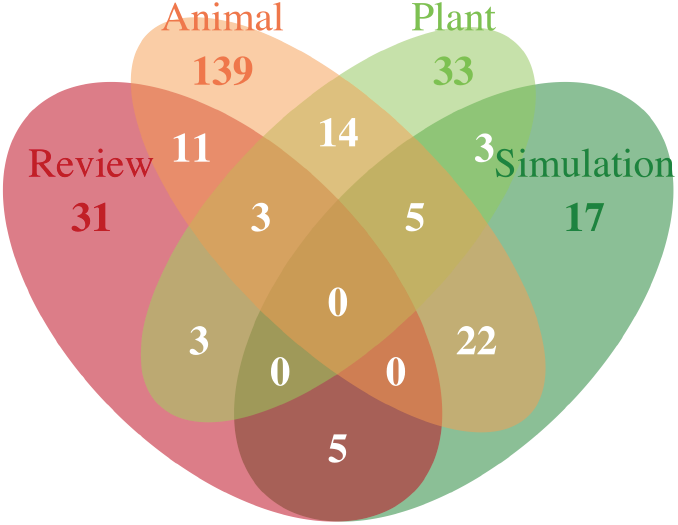
Classification of manuscripts self-identified as 'landscape genetics’ into *Review, Simulation, Animal*, and *Plant* categories (see Table 1 for a complete description).

### Fusion Analysis

The discriminant functions were relatively well behaved with a total probability of misclassification (*e.g.*, the observed probability of classifying an LE manuscript as PG or *vice versa)* of *TPM*_*Intro*_ = 0.025 for the *Introduction* sections. The classification of LG papers based upon the *Introduction* were unequal with a relative frequency of classification to PG over LE of 2:1 (*p*_*PG*_=0.65 vs. *p*_*LE*_= 1 - *P*_*pg*_ = 0.35), a ratio that deviates significantly from even allocation to both progenitor disciplines (binomial test, x=91, n=139, P=3.33e^-4^, Cl_95_= 0.57-0.73). The analysis of LG *Methods* sections showed both a lower error rate for classification (*TPM*_*Methods*_ = 0.020) and a more pronounced bias, a 3:1 ratio, to classify LG papers as PG (*p*_*PG*_=0.81) instead of LE (*p*_*LE*_=0.19; binomial test for equal allocation, x=112, n=139, P=1.68e^-13^, Cl_95_= 0.73 - 0.87). As a consequence, research manuscripts identified as landscape genetic are much more similar to manuscripts classified as population genetics than to manuscripts identified as landscape ecology. Density estimation for LE, LG, and PG discriminant scores from these analyses shows the extent to which LG is intermediary, though highly skewed, between the distribution defining LE and PG manuscripts (Figure 2).

**Figure 2.**
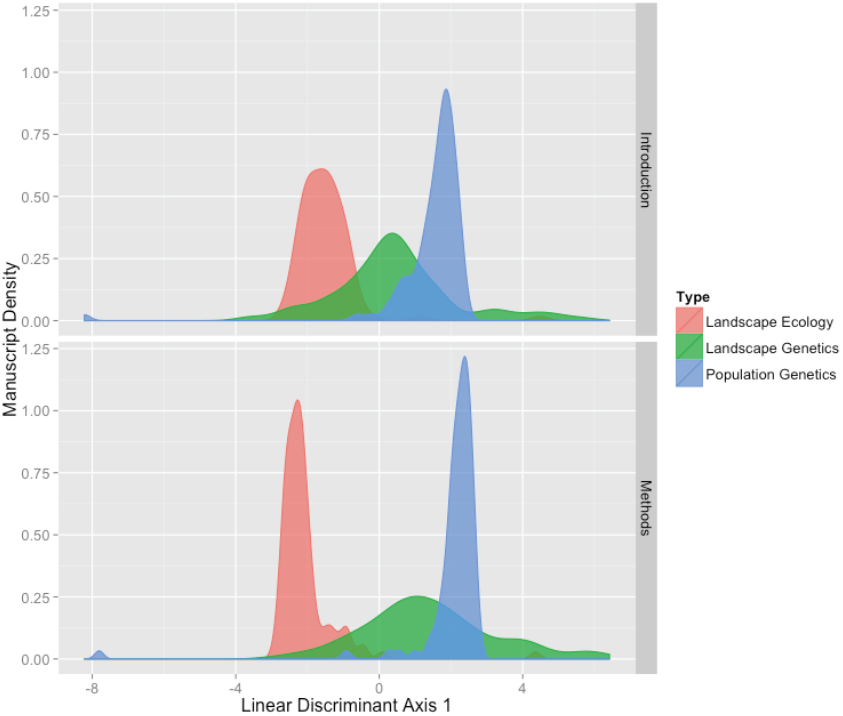
Density plot of discriminant scores for LE, LG, and PG papers defined by the contents of the *Introduction* (top) and *Methods* (bottom) sections. The majority of LG papers were classified as PG for both the *Introduction* (65%) and *Methods* (78%) textual components.

### Circumscribing Landscape Genetics

Differences in term usage between LG, LE, and PG papers (Table 2) highlight the particular focus of landscape genetic studies and how they differ from LE and PG manuscripts. In the *Introduction* section, the specific word stems *distanc, barrier*, and *connect* were all found to be overrepresented in LG manuscripts. That is not to say that these terms were not found in LE or PG manuscripts, in fact, they occurred at a rate of 0.21, 0.09, and 0.20 for LE papers and 0.16, 0.18, and 0.15 for PG papers (respectively), they were just more commonly found in LG manuscripts. The *Methods* sections showed more idiosyncratic differences with only one term found at elevated frequencies in LG manuscripts over both LE and PG: *mantel.* It is perhaps not surprising given the overrepresentation of *distanc*, that *mantel* (the most common way in which distance matrices are evaluated) was found in the *Methods* section. As in the *Introduction* section, none of the most overrepresented terms in LG papers were missing from the other manuscripts’ categories (LE frequencies range for top LG terms: 0.01-0.41; PG frequency range: 0.03-0.53), they were just used at elevated frequencies.

**Table 2.** Differences in the frequency of stemmed terms comparing landscape genetic (LG) to both Landscape Ecology (LE) and Population Genetic (PG) manuscripts in both (A) *Introduction*, and (B) *Methods* sections. Numerical value next to stemmed term represents the frequency difference in term occurrence across manuscripts.

| Landscape Ecology |  |  |  | Population Genetics |  |  |  |
| --- | --- | --- | --- | --- | --- | --- | --- |
| +LG |  | +LE |  | +LG |  | +PG |  |
| genet | 0.89 | veget | 0.34 | landscap | 0.94 | evolut | 0.37 |
| gene | 0.88 | resourc | 0.26 | spatial | 0.58 | genom | 0.30 |
| barrier | 0.53 | properti | 0.24 | featur | 0.51 | sequenc | 0.27 |
| dispers | 0.53 | densiti | 0.24 | distanc | 0.51 | morpholog | 0.27 |
| differenti | 0.52 | soil | 0.24 | connect | 0.50 | mutat | 0.23 |
| flow | 0.48 | activ | 0.23 | habitat | 0.50 | histori | 0.21 |
| geograph | 0.47 | abund | 0.22 | structur | 0.48 | trait | 0.21 |
| distanc | 0.46 | communiti | 0.22 | dispers | 0.47 | origin | 0.17 |
| connect | 0.45 | control | 0.21 | barrier | 0.44 | genus | 0.16 |
| isol | 0.44 | matter | 0.20 | movement | 0.41 | fitness | 0.16 |

Table 2 (cont.) (B) Methods
| Landscape Ecology |  |  |  | Population Genetics |  |  |  |
| --- | --- | --- | --- | --- | --- | --- | --- |
| +LG |  | +LE |  | +LG |  | +PG |  |
| genotyp | 0.83 | day | 0.33 | landscap | 0.80 | align | 0.34 |
| pairwis | 0.81 | veget | 0.32 | spatial | 0.62 | substitut | 0.31 |
| gene | 0.81 | open | 0.32 | mantel | 0.55 | phylogenet | 0.26 |
| dna | 0.79 | survey | 0.30 | area | 0.48 | neutral | 0.19 |
| mantel | 0.78 | soil | 0.29 | habitat | 0.48 | evolut | 0.19 |
| differenti | 0.78 | land | 0.29 | dispers | 0.44 | segreg | 0.19 |
| equilibr | 0.77 | normal | 0.28 | barrier | 0.44 | recombin | 0.18 |
| linkag | 0.77 | domin | 0.28 | connect | 0.43 | experi | 0.18 |
| pcr | 0.76 | composit | 0.27 | correl | 0.43 | phenotyp | 0.17 |
| popul | 0.73 | annual | 0.26 | distanc | 0.43 | ident | 0.16 |

A hierarchical analysis of the *Methods* section for self-identified LG papers showed clear clustering of manuscripts (Figure 3). Bootstrapping the stemmed terms revealed a total 13 separate clusters of manuscripts whose nodes were supported with at least 95% confidence. The main bifurcation in Figure 3 is just shy of that at 94% bootstrap support and is indicated as shown. It appears that as a group, there is not a single kind of manuscript that characterizes landscape genetics but rather there are divergent categories. To aid in summarizing the differences between the main groups, the relative frequency of stemmed term usage was estimated to identify overrepresentation of particular terms unique to that group and is depicted by wordcloud inserts for the top ten most overrepresented terms in each cluster. The smallest cluster has only three manuscripts and the relative frequency of term usage should be interpreted with extreme caution, the others have 84 (lower clade) and 52 (upper clade) manuscripts providing a more robust estimation of relative frequency bias. The intersection of term usage in the two large clades reveal the stemmed terms *test, distanc, calcul, estim*, and *data* in common. More interesting are the terms not shared among the top ten terms. The terms in the upper clade (in decreasing order of usage) include *individu, model, valu, base*, and *compar*, whereas the unique terms for the lower clade are *sampl, popul, number, studi*, and *analysis.* The differences seen do not represent words unique to the clade, rather they occur at an increased frequency when compared to the remaining manuscripts. Of note, and particularly critical to the differences as depicted by term usage, is that the upper clade emphasis on *individu* and the lower one that utilizes the term *popul*, emphasizing potentially divergent foci for research questions. The remaining terms overrepresented in each clade were much more generic including *model, valu, base, compar, sampl, number, studi*, and *analysis.*

**Figure 3.**
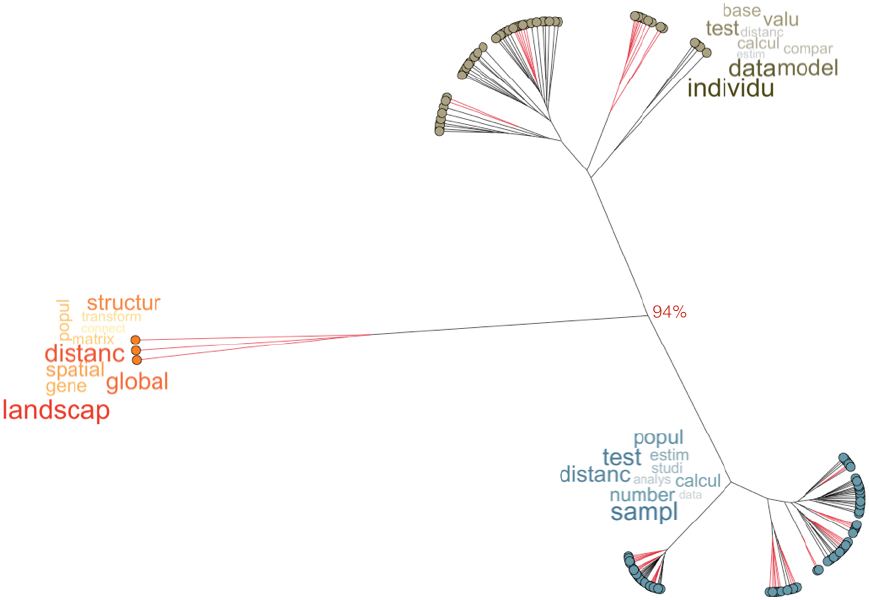
Hierarchical clustering using Ward's minimum variance metric of stemmed text content found within the *Methods* section of self-identified landscape genetic papers. Clusters with >95% bootstrap support are shown with red branches (the main bifurcation separating the three groups had 94% bootstrap support as indicated). The relative frequency of to the top 10 stemmed words are shown as wordcloud inserts (term font size represents decreasing order of usage).

## Discussion

As a whole, the data show that self-identified landscape genetic papers overwhelmingly resemble studies in population genetics, both in how the questions are formulated (a 2:1 allocation bias) and the methodologies that are used to answer them (a 3:1 bias). The relative term frequency usage for both the *Introduction* and *Methods* sections (Table 2) suggest that as a whole, both landscape ecology and population genetics are much broader fields addressing a wider array of question as indicated by the the maximum frequency bias for both landscape ecology and population genetic studies being lower than the top ten terms defining landscape genetics. The differences between landscape ecology and landscape genetics are denoted by the use of terms such as *genet, gene, genotyp, dna, linkag*, and *pcr*, clearly related to the addition of genotypic data and highlighting the ease at which the two fields may be differentiated. That is not to say that the history of landscape ecology is entirely devoid of the use of population genetic theory (e.g., Manicacci *et al.* 1991), the distinction simply provides an identifiable delineation. When compared to population genetics, terms such as *landscap, spatial, barrier*, and *distanc* are overrepresented in landscape genetic studies. As previously noted, it would be a fragile argument to suggest that population genetic studies are distinct from landscape genetic ones because they do not focus on spatial structure, barriers, or the use of distance measures, these were all present in population genetics well before this new moniker was established. Overall, both the discriminant analysis and term usage suggest that to a large degree landscape genetic studies resemble the kinds of questions and approaches commonly called population genetic prior to 2003. These data do suggest that there should be a discussion about the appropriateness of referring to landscape genetics as a unique endeavour versus a subdiscipline residing within population genetics.

An additionally interesting set of bifurcations is found within the totality of landscape genetic studies themselves. As a group, landscape genetics is dominated by animal systems, outnumbering research projects focusing on plant systems or theory & simulation as well as general review articles (of which there are indeed many) combined (Figure 1). It is at least in part due to the overrepresentation of animal-focused studies that the methodologies used for landscape genetics shows deep bifurcation (Figure 3), particularly as they apply to the distinction between the clade identified by the overrepresentation of *individu* and the one using *popul* more frequently. It has been argued that logistical issues associated with life history traits of some kinds of species such as bears (Cushman *et al.* 2006), moose (Finnegan *et al.* 2012), pika (Castillo *et al.* 2014), and tortoise (Hagerty *et al.* 2011) prevent population-level sampling schemes. It is not to say that certain species cannot be examined at the population level, rather at a more fundamental level, the kinds of *questions* being asked in these systems are different than those being applied to organisms that are examined within a population level context.

Individual-based studies often use genetic markers as a tool to understand transient individual movement across heterogeneous landscapes, particularly focusing on how it is mitigated through individual behavioral decisions. For example, in the course of introducing the utility of a Causal Modeling approach, Cushman *et al.* (2006) showed that movement costs based upon inter-individual genetic distances in black bears were most highly correlated with landscape gradients in land cover along specific elevations. They conclude that gene flow, as estimated by individual movement patterns across the sampled landscape, is most common through regions at elevations of ~1,000m and through stands of continuous forest canopy. While these findings provide concrete data on demographic movement and aid in the development of conservation and management practices, the authors correctly warn that there is not a direct link between these individual behavioral decisions and actual gene flow, as it is characterized as a microevolutionary process and as population genetic theory defines it. More recently, a study in caribou by Yannic *et al.* (2013) showed that the strength of the correlation between inter-individual relatedness and ecological separation, a study of isolation by ecological distance, varied dramatically across both year and the sex of the individual being followed. These correlations peaked during calving and rutting periods (maximum Mantel r = -0.25) and dropped throughout the rest of the year. Even if there are correlations between relatedness and ecological separation fluctuating through time, there does not yet appear to be a theoretical foundation through which quantitative measures gained from individual-based landscape genetic analyses are connected to parameters relevant to evolutionary structure and stability such as F_ST_ and N_e_.

Conversely, approaches that use population-based sampling in landscape genetics do provide a more direct linkage between heterogeneity in the landscape and population genetic structure. This connection is made possible because the bulk of existing microevolutionary theory, what we call population genetics dating from Cotterman (1940), Wright (1943) and Malécot (1948), is based upon population-level inferences. As a result, the parameters estimated to describe inter-population differentiation, either genetic distance or structure based, have a more direct connection to existing evolutionary theory. While most of the individual based studies are in animals, population-level studies are being applied across all kinds of life histories and sampling approaches. Examples include organisms such as pond-dwelling amphibians (e.g., Murphy *et al.* 2010, Moore *et al.* 2011), or among plant groups such as herbs and succulents (e.g., Dyer *et al.* 2010, Matter *et al.* 2013), understory trees (e.g., Dyer *et al.* 2012, DiLeo *et al.* 2014), or canopy trees (e.g., Diniz-Filho *et al.* 2009, Poelchau & Hamrick 2012, Wei *et al.* 2013).

A direct connection between individual processes is largely missing from metrics based upon inter-individual measurements (though see McRae 2006 who argues that ecological separation approximates a coalescent process). Perhaps in response to this, a large portion of empirical landscape genetic studies rely upon stochastic simulation approaches (e.g., Figure 1) as a way to circumvent, either consciously or otherwise, the lack of theoretical connections. The benefit of a simulation-based approach is that one can examine highly specialized situations, though this benefit is paid for by a lack of generality. Simulation alone cannot take the place of theory, a criticism that has been leveled in population genetics as well; running *ms* is not the same as deriving closed form solutions to population genetic problems. In time, perhaps individual-based approaches can be reinforced by the development of population genetic theory providing a stronger connection between how transient individual movement translates into quantifiable measurements on processes influencing population genetic structure. However at present, despite the suggestions by Manel *et al.* (2003), there does not appear to be a robust theoretical basis for how landscape genetics is different from existing bodies of knowledge.

Despite the differences between individual- and population-based approaches to landscape genetics, the overlap in the methodologies highlight potentially the main contribution of landscape genetics to date, namely the use of distance as a primary analysis tool. Landscape genetic papers use the stemmed term *distanc* much more frequently in setting up research questions (+46% over landscape ecology and +51% over population genetics; Table 2). A similar level of overrepresentation was found in the *Methods* sections when compared to the sample of population genetic manuscripts (+43%).

The key distinction here is that the increased use of distance approaches as a paradigm for analyses has grown tremendously. Distance matrices are used for both predictor (ecological, spatial, topographic) and response (observed genetic data) variables. Mechanistically, there is a wide array of approaches available to quantify the predictor variables, often encoded as spatially explicit raster data, including least-cost paths (e.g., Walker & Craighead 1997, Graham 2001, Adriaensen 2003), corridor connections (Epps *et al.* 2007), and all paths (e.g., circuit theory, McRae 2006), to estimate an among-site (or individual as the case may be) distance matrix. As most things in Biology, in certain situations one of these approaches can provide a better fit to the observed genetic data than the others, though all too often alternatives are not tested and one is used based upon supposed ‘expert opinion.’ Depending upon the kind of raster representation, a suite of new terms such as isolation by landscape resistance (denoted IBR; McRae 2006), isolation by habitat (IBH; Mallet *et al.* 2014), isolation by ecology (IBE; Wang *et al.* 2012, Shafer & Wolf 2013), and isolation by barriers (IBB; Cushman *et al.* 2006) has sprung into existence. Despite the perceived importance that alternative raster data has, the realized differences in these IB* models are minor compared to the similarity in how we use them. These are essentially models based upon the classic isolation by distance (IBD; Wright 1943) paradigm, which had previously been applied to genetic data using spatial (Euclidean) separation. Independent of the differential connection to theory between individual and population level approaches, it is not entirely clear if the proliferation of IB* models is warranted for more than keyword inflation as they all answer the same general research question. In reality the forces that have produced the observed spatial distribution of genetic structure are most likely due to the interaction of several of these processes overlain through time (e.g., Dyer *et al.* 2010). It remains to be seen if the application of general IB* frameworks across a broader range of studies results in a reticulation of terminology back to generalized IBD models or progresses towards continued lexicographic fragmentation.

### Looking Forward

At a base level, the general model tested in majority of landscape genetic studies can be quantified as:

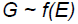

where *G* is some measure of genetic composition such as pairwise *F*_*ST*_ (Rousset 1997; or one of its many analogs), multilocus divergence (Smouse *et al.* 2001), or one of a variety of potential estimates of genetic frequency (e.g., Weir 1990) or genetic network (Dyer *et al.* 2010) distance. This response is fit using some model (e.g., the *~f()* part) to the set of predictor variables describing a subset of ecological and/or spatial separation. The variables represented by *E* can include highly detailed GIS layers, rough approximations of bioclimatic conditions, physical separation based upon distance or barriers, or it can just be populations as a categorical variable.

The way in which both *G* and *E* are analyzed in landscape genetic studies is most commonly conducted using either correlative approaches such as the single Mantel or Partial Mantel tests (though see Legendre & Fortin 2010) or via non-parametric regression approaches (Lichstein 2007). The salient distinction of which is that the latter can produce a predicted genetic distance surface (e.g., Dyer *et al.* 2010, specifically Figure 4 therein) suitable for subsequent hypothesis derivation and testing whereas the former cannot.

Comparatively, the level of sophistication that has been developed to create the components of this relationship (i.e., *G, E*, and *~f()*) is highly skewed. Largely borrowed from landscape ecology, the characterization of physical and ecological separation between observations as depicted in *E* is highly developed and rather sophisticated. The various IB* predictors and the models within which they are analyzed such as least cost path and circuit theory, are continuing to be developed, are quite novel, and very precise. These developments offer the researcher a wide range of potential predictor variables and variable configurations. To a lesser degree, the statistical approaches based upon the analysis of distance matrices have seen similar development and refinement. Largely, work on understanding how to conduct the analyses has focused on refining the tools we already have; a better Mantel, a more sophisticated Moran’s I, etc. The consequence of this is that we’ve been able to ask the same kinds of questions in a more precise manner, though we are still asking the same questions.

Unfortunately, the way in which we characterize *G* has generally languished. The use of pairwise genetic structure is an aged approach. The use of pairwise structure may be quite robust in the long term but is often too slow to track contemporary processes (e.g., Dyer 2007). Moreover, it often fails to capture the peculiar nature of how contemporary microevolutionary processes such as heterospecific interference in mating (Dyer & Sork 2001), asymmetry in dispersal (Papa & Gept 2003), and contemporary consequences of fragmentation (Fore *et al*. 1992) influence genetic variation unless the signal is rather strong (Jaquiéry *et al.* 2011). Many pairwise genetic distance metrics also come with a set of deep time assumptions (e.g., Roger’s, Nei’s, and Bhattacharya’s) or have relatively little theory supporting their connection to microevolutionary processes (e.g., Bray-Curtis distance). Other population genetic parameters, less often encountered in landscape genetic studies, such as inbreeding (Diniz-Filho *et al.* 2009, Guarino & Cipriani 2013), relatedness or coancestry (Waterhouse *et al.* 2014), and effective population size (Wang *et al.* 2011, McCracken *et al.* 2013) have the potential to provide direct links between landscape processes and established evolutionary theory, though they are often overlooked. One novel approach to characterizing the *G* side of the equation, distinct from the estimate of pairwise approaches, is the development of network approaches based upon conditional genetic covariance (Dyer & Nason 2004, Dyer *et al.* 2010, 2012) where differences are estimated utilizing the entirety of the data set rather than set of pairwise contrasts. While these approaches have allowed the movement beyond classical population genetic parameters, active development in the way in which we characterize population genetic data to bring it to the level at which landscape ecologists have developed spatial data presentation is sorely needed. It is time for the population geneticists to become more involved in landscape genetics. There is a real opportunity here to not only introduce existing tools that are not being utilized to their full potential, but also to push the envelope in how we characterize genetic data and how we connect landscape scale processes (both spatial and temporal) into the existing body of evolutionary theory.

## Conclusion

After looking at how landscape genetic studies are portrayed through a textual analysis of published manuscripts, it appears that the field is neither unique nor uniform. It is borrowing disproportionately from population genetics and to a lesser extent landscape ecology. There does not seem to be an indication that landscape genetics addresses questions and hypotheses that other disciplines cannot answer despite the initial suggestion that it does so, relegating it to perhaps more multidisciplinary in nature than interdisciplinary.

A reviewer of this manuscript posited that just because the data presented herein does not support the notion that landscape genetics has yielded insights unattainable by either landscape ecology or population genetics, it does not mean that it will not. Indeed, the purpose of this opinion piece is to evaluate the interdisciplinary characteristics of landscape genetics and to look at the body of work in an alternative way with the hopes of finding the components that need additional development. These data shed some light on some of the things that need further development of landscape genetics to live up to what Manel *et al.* (2003) originally envisioned—undoubtedly more exist.

At a more base level, what we call the kind of science we do may or may not be relevant and this may all be about semantics that at the end of the day are immaterial. My impetus here is to not argue semantics but to understand where additional developments may yield the most reward. As a self-identifying body of research, landscape genetics appears to be currently fragmented. The data presented suggest that there are at least two broad groups roughly partitioned by the focus on individual versus population level processes, both of which seek to address somewhat overlapping sets of hypotheses.

Looking forward, there are several potential avenues available to landscape genetics as it continues to mature. First, it could continue on its present course representing a loosely confederated group of different individual and population level studies examining the consequences of heterogeneous landscapes using distance matrices and Mantel tests. This is probably the least beneficial route as it essentially leaves landscape genetics as a grab bag of loosely related questions and approaches. It is also possible that one of the groups may be subsumed under a yet to be coined moniker, something perhaps all too common in biological sciences, though allowing both sections (individual transient movement vs. effects landscapes have on population genetic structure) to focus more intently on how the specific kinds of hypotheses these approaches provide broad biological relevance. Lastly, and the route for which I advocate, is that the relevant theory connecting individual and population level analyses be developed. While landscape genetics has generally been using population genetic hypotheses and methodologies, it has only been using a small fraction of even the available tools and is ripe ground for continued theoretical development. Independent of which trajectory the field takes, the continued integration of spatial and ecological contexts into the analyses of microevolutionary processes will only aid in tightening the connections between the fields of Ecology and Evolution, *sensu lato*, and perhaps more importantly spark continued refinement of the kinds of questions we address.

## Acknowledgements

This manuscript was adapted from an invited seminar arranged by N. Balkenhol. I also thank A. Eckert, B. Verrelli, and H. Wagner for ongoing discussions on the nature of Landscape Genetics and its relationship to both Landscape Ecology and Population Genetics.

## Data Accessibility

The data and scripts necessary to perform all analyses in this manuscript are posted to github. The R library footprint is located at https://github.com/dyerlab/footprints and the scripts and data for this paper hosted at https://github.com/dyerlab/ITSATALG. Both components are available under an open source license.

